# Development of high-throughput screening assays for profiling snake venom Phospholipase A_2_ activity after high-resolution chromatographic fractionation

**DOI:** 10.1101/2020.01.20.912758

**Authors:** Kristina B.M. Still, Julien Slagboom, Sarah Kidwai, Chunfang Xie, Bastiaan Eisses, Freek J. Vonk, Govert W. Somsen, Nicholas R. Casewell, Jeroen Kool

## Abstract

Many organisms, ranging from plants to mammals, contain phospholipase A_2_ enzymes (PLA_2_s), which catalyze the production of lysophospholipids and fatty acid proinflammatory mediators. PLA_2_s are also common constituents of animal venoms, including bees, scorpions and snakes, and they cause a wide variety of toxic effects including neuro-, myo-, cyto-, and cardio-toxicity, anticoagulation and edema. The aim of this study was to develop a generic method for profiling enzymatically active PLA_2_s in snake venoms after chromatographic separation. For this, low-volume high-throughput assays for assessment of enzymatic PLA_2_ activity were evaluated and optimized. Subsequently, the assays were incorporated into a nanofractionation platform that combines high-resolution fractionation of crude venoms by liquid chromatography (LC) with bioassaying in 384-well plate format, and parallel mass spectrometric (MS) detection for toxin identification. The miniaturized assays developed are based on absorbance or fluorescence detection (respectively, using cresol red or fluorescein as pH indicators) to monitor the pH drop associated with free fatty acid formation by enzymatically active PLA_2_s. The methodology was demonstrated for assessment of PLA_2_ activity profiles of venoms from the snake species *Bothrops asper*, *Echis carinatus*, *Echis coloratus, Echis ocellatus*, *Oxyuranus scutellatus* and *Daboia russelii russelii*.

## 1. Introduction

Phospholipase A_2_ (PLA_2,_ EC 3.1.1.4) is a common enzyme occurring in many organisms, ranging from plants to mammals.[1] PLA_2_s are crucial enzymes in lipid metabolism and lipid-protein interactions. They also play a significant role in various cellular processes such as biotransformation and digestion of phospholipids, signal transduction and host defense.[2] PLA_2_s are connected to human pathophysiological events and associated with certain types of cancers, arthritis, and inflammatory disorders, and therefore have been studied extensively.[3–5]

PLA_2_s are also widely distributed in animal venoms. Snake venom PLA_2_s (svPLA_2_s) can be major toxin components – they often comprise 30-71% of the total venom proteins, and they can also be diverse in terms of amino acid composition, as evidenced by the UniProtKB database containing over 400 unique svPLA_2_s.[6] In addition to catalyzing the production of lysophospholipids and fatty acid pro-inflammatory mediators, svPLA_2_s are multifunctional enzymes and cause a wide variety of toxic effects ranging from neurotoxicity, myotoxicity, anticoagulant effects, cytotoxicity, cardiotoxicity, to edema.[7] Because of the pathological consequences of these toxins to prey and snakebite victims, svPLA_2_s have been extensively investigated.[7, 8] Since May 2018, snake envenoming has been categorized as one of the most neglected tropical diseases.[9] The annual number of human snakebites is estimated to be between 1.8 and 2.7 million worldwide, resulting in 81,000–138,000 deaths and around three times more cases of permanent morbidity cases.[10] Many victims receive inadequate treatment or no treatment at all. The World Health Organization (WHO) has therefore adopted a resolution towards tackling this devastating global health problem.[10]

Snake venoms are complex biochemical mixtures that are injected via the fangs of snakes for killing prey and for defense. These venoms comprise many different pathological proteins and peptides, and other organic molecules. Venom composition is typically complex (50-200 proteins per species) and highly variable, with often extensive inter-specific, and even, intra-specific venom variations observed.[11, 12] Rooted in the latter are geographical differences, living habitat, sex, and age of a snake, thereby increasing venom complexity even more.[13, 14] Consequently, antivenom therapies, which are based on antibodies produced in horses or sheep following their immunization with venom or mixtures of venoms, are highly specific to those venoms used for production, but often lack efficacy for treating snakebites by other snake species.[12] In recent years, research efforts have increased on the potential utility of small molecule drugs as potential alternatives for conventional antivenom therapies. These promising approaches focus on neutralizing entire classes of venom enzymes, irrespective of sequence and structural variation, with small molecular inhibitors or metal chelators. For instance, recent studies have demonstrated that enzyme inhibitors and metal chelating agents, such as EDTA and DMPS, are capable of neutralizing snake venom metalloproteinase toxins in pre-clinical models of envenoming.[15, 16] In relation to svPLA_2_s, a number of studies have explored the neutralizing potential of the generic PLA_2_ inhibitors methyl-varespladib and varespladib.[17] These compounds show great promise for the future development of affordable, stable and broad-spectrum treatment for PLA_2_-induced toxicities following snake envenoming. However, as svPLA_2_s are major venom toxins responsible for a diverse array of severe pathologies following envenoming, the development of analytics for rapid svPLA_2_ profiling after chromatographic separation of snake venoms – the approach described in this study - is an essential prerequisite for efficient selection and *in vitro* validation of novel PLA_2_ inhibiting agents.

Assays using chromogenic molecules for PLA_2_ profiling have previously been developed.[18, 19] One of these assays relies on hydrolysis of a substrate by PLA_2_s into a coloured product, with ensuing absorbance measured at 425 nm, as described by Petrovic *et al*.[18] In addition, both radioactivity based assays [20, 21] and fluorogenic assays [22, 23] have been described. All these assay formats, however, are probe substrate dependent and thus dependent on the selectivity and enzymatic activity of each PLA_2_ for the probe substrate used. Assays for assessment of generic enzymatic PLA_2_ activity use a phospholipid as a generic substrate, in combination with pH indicators to measure medium acidification upon phospholipid hydrolysis.[24–26] This enzymatic reaction involves PLA_2_s cleaving the sn-2 ester of phospholipids, generating a free fatty acid and a lyso-phosholipid. Price *et al*. and Lobo De Araljio *et al*. measured PLA_2_ activity of crude snake venoms using the pH indicators bromothymol blue [26] and phenol red [27], respectively. These assays were performed in cuvettes, using a spectrophotometer, or in 96-well format. The use of bromothymol blue in a PLA_2_ assay was considered for this study, but was abandoned since literature reports this pH indicator to be able to inhibit phospholipase subunits under certain conditions.[27]

In this study, two assays for enzymatic PLA_2_ activity were developed in 384-well format. One assay makes use of cresol red with colorimetric readout whereas the other uses fluorescein with fluorescence readout. The assays were optimized and validated for application in a workflow comprising high-resolution chromatographic fractionation of snake venoms followed by enzymatic PLA_2_ bioassaying. For this, well plates with collected fractions were vacuum-centrifuged to dryness followed by robotic pipetting of bioassay reagents and plate reader based readout in order to assess PLA_2_ activities. The feasibility and usefulness of the approach for measuring generic svPLA_2_ activity profiles was demonstrated using medically relevant snake venoms from *Bothrops asper*, *Echis carinatus*, *Echis coloratus, Echis ocellatus*, *Oxyuranus scutellatus* and *Daboia russelii russelii*.

## 2. Materials and Methods

### 2.1 Chemicals and biological reagents

Water was purified with a Milli-Q Plus system (Millipore, Amsterdam, The Netherlands). DMSO was supplied by Riedel-de-Haën (Zwijndrecht, The Netherlands). Acetonitrile (ACN; ULC/MS grade) and formic acid (FA) were obtained from Biosolve (Valkenswaard, The Netherlands). All salts used for buffer preparation were of analytical grade and purchased from Merck (Kenilworth, USA), Fluka (Bucharest, Romania) or Sigma-Aldrich (Darmstadt, Germany). Micro-90^®^ concentrated cleaning solution was supplied by Sigma-Aldrich. Lyophilised snake venoms (see Table 1) were provided by the Centre for Snakebite Research & Interventions (Liverpool School of Tropical Medicine, UK) and stored long-term at −80 °C. Stock solutions of crude venoms (5.0 mg/mL) were prepared in water prior to analysis and stored at −80 °C. A 1 mM Tris (pH 8) buffer solution for the bioassay was made in Milli-Q water and its pH was checked at room temperature. After preparation, the buffer was stored at 4 °C until use. Cresol red and fluorescein were from Sigma-Aldrich. The 5 mM cresol red stock solution was prepared in methanol and stored at −20 °C. The 1 mM fluorescein stock solution was prepared in DMSO and kept at −80 °C. Triton-X-100 was purchased from Thermo Scientific (Landsmeer, Netherlands) and a stock solution of 170 mM was prepared in milli-Q water and stored at −20 °C. Phosphatidylcholine from soy beans was purchased from Sigma-Aldrich of which a stock solution of 20 mg/mL was prepared in methanol and kept at −20 °C.

**Table 1.** List of analyzed snake venoms and their abbreviations as used in this study.

| Snake venom | Origin | Abbreviation |
| --- | --- | --- |
| *Bothrops asper* | Costa Rica | BA |
| *Echis carinatus* | India* | EC |
| *Echis coloratus* | Egypt | ECO |
| *Echis ocellatus* | Nigeria | EO |
| *Oxyuranus scutellatus* | Australia | OS |
| *Daboia russelii russelii* | Sri Lanka | DRR |

\* Note that the Indian *E. carinatus* venom was collected from a single specimen that was inadvertently imported to the UK via a boat shipment of stone, and then rehoused at LSTM on the request of the UK RSPCA.

### 2.2 PLA_2_ assay using cresol red as pH indicator

Pre-prepared 1.0 mM TRIS buffer solution was used at room temperature and pH 8.0. The assay was performed with all other reagents at room temperature, which is crucial due to the pH dependence of the assay. For best performance the assay reagent mix was freshly prepared in a 50 mL PP Centrifuge tube (Corning Life Sciences B.V., Amsterdam, The Netherlands). The mix contained NaCl (75 mM), KCl (75 mM), CaCl_2_ (7.5 mM), Cresol Red (0.037 mM), Triton-X-100 (0.66 mM) and phosphatidylcholine (0.66 mM) in 1.0 mM Tris (pH 8.0). The salts were added as dry compounds (after accurate weighing), whereas for the other constituents, accurate volumes of the stock-solutions (Section 2.1) were used. Triton-X-100 is needed to increase the solubility of the substrate, improving the interaction between PLA_2_ and phosphatidylcholine. Direct addition of Triton-X-100 resulted in homogeneity problems and unsatisfactory assay performances. Therefore, the use of a pre-prepared stock solution of Triton-X-100 (170 mM) was used. The phosphatidylcholine solution was added last, as it slowly degrades upon contact with water. Prior to addition of phosphatidylcholine, the pH of the total solution was always checked and, if needed, adjusted to pH 8.0. The buffer capacity of the assay solution was low in order to allow measurement of a pH drop as result of PLA_2_ activity. The assay was initiated by robotically pipetting 40 µL of the final assay reagent mix into each plate well containing either vacuum centrifuge-dried snake venom fractions or 10 µL of test solution used for assay development. In the latter case, concentrations of the assay mix constituents were adjusted to match the final assay concentrations stated above. The plate was placed in the plate reader within 5 min after pipetting, and the plate reader was thermostated at 25 °C. The absorbance of each well content was measured at 572 nm with a Thermo Fisher Scientific Laboratory Varioskan™ LUX Multimode Microplate Reader using SkanIt 4.1 (Landsmeer, Netherlands). Measurements were performed in one kinetic loop with a total measurement time of 40 min. Two data-processing options in the SkanIt 4.1 software were used to determine PLA_2_ activity from the measured kinetic curves: (1) slope of a reading range (for well plates holding venom fractions), and (2) average rate in time per well (during assay development). For the latter, one measurement data point was plotted every 10 min over the total measuring curve. For constructing bioassay chromatograms of snake venoms, for each well the value resulting from the processed assay data was plotted against the LC elution time corresponding to the well.

### 2.3 PLA_2_ assay using fluorescein as pH indicator

In this PLA_2_ activity assay fluorescein is used as a fluorescent pH indicator in black 384-well, F-shape microplates (Greiner Bio One, Alphen aan den Rijn, the Netherlands). A decrease in assay pH causes a decrease of fluorescence intensity. Concentrations and preparation of the assay reagents and mix were the same as for the colorimetric assay (Section 2.2), but no Cresol Red was added. Instead, fluorescein was present at a final concentration of 1 µM. The assay was started by using the same procedure as stated in Section 2.2. The plate reader temperature was set at 25 °C and of each well fluorescence was measured using an excitation wavelength of 488 nm and an emission wavelength of 520 nm in the Varioskan™ LUX Multimode Microplate Reader. Measurements were performed in one kinetic loop with a total measurement time of 40 min. The bioassay data were presented as bioassay chromatograms when used for snake venom screening.

The cleaning procedure of the robotic pipetting machine is as reported previously in Still et al.[28] For typical kinetic curve results obtained when running both PLA_2_ assay formats with venoms high in PLA_2_ abundance, the reader is referred to supporting information S1. The difference in curve slopes allows for the reconstruction of bioactivity chromatograms (more details in Section 3.3).

### 2.4. Instrumental setup for venom fractionation

For high-resolution fractionation of snake venoms, the analytical system previously described by Still *et al*.[28] and by Mladic *et al*.[29] was used. Samples were injected with a Shimadzu SIL-30AC autosampler and LC separation was performed with a Shimadzu LC system controlled by Lab Solutions software. Gradient LC was performed using two Shimadzu LC-30AD pumps (A and B) operated at a total flow rate of 0.5 mL/min. Mobile phase A was water–ACN–FA (98:2:0.1, v/v/v) and mobile phase B was water–ACN–FA (2:98:0.1, v/v/v). The gradient was as follows: 0% to 50% B (20 min), 50% to 90% B (4 min), 90% B (5 min), 90% to 0% B (1 min), 0% B (10 min). A 150 × 4.6 mm ID analytical column packed with Xbridge^TM^ BEH300 reversed-phase C18 material (5 µm) was used for separations and was maintained at 37 °C in a Shimadzu CTD-30A column oven. The column effluent was split in a 1:9 ratio using a low-dead-volume flow splitter. The flow of 0.05 mL/min was either directed to waste or led to a high-resolution time-of-flight (TOF) mass spectrometer for compound identification. During assay and method development, no MS data was acquired. The flow of 0.45 ml/min was led to a either a Gilson 235 robot programmed as fractionation device or a FractioMate^TM^ fractionator (SPARK-Holland & VU, Netherlands, Emmen & Amsterdam) each providing fractions (6 s/well) onto clear or black 384-well plates. Fractionation was controlled by employing in-house written Ariadne software or FractioMator software (Spark-Holland & VU), respectively. The plates with fractions were vacuum centrifuged to dryness at room temperature using a Christ Rotational Vacuum Concentrator RVC 2-33 CD plus (Salm en Kipp, Breukelen, the Netherlands) with a cooling trap at −80 °C, and then stored at −80 °C.

## 3. Results and Discussion

This research focused on the development and optimization of enzymatic PLA_2_ activity assays suitable for application in the nanofractionation platform to allow profiling of venom fractions. For this, these assays should be sensitive, rapid, robust, and applicable to small LC fractions (collected in 384-well plate format). PLA_2_ catalyzes the hydrolysis of ester bonds of glycerophospholipids at position sn-2. PLA_2_ activity assaying can be based on the accompanying acidification of the reaction mixture caused by the formed free fatty acids during phosphatidylcholine hydrolysis. In presence of pH indicators, activity will cause measurable color changes. The use of bromothymol blue in the required miniaturized assay was considered for use in this study, however, (i) the literature reports this pH indicator to inhibit phospholipase subunits, [26] while pilot experiments using bromothymol blue for assaying venoms showed highly varying and non-repeatable results. Specifically, our pilot results appeared unrelated to PLA_2_ hydrolysis of phosphatidylcholine, and therefore bromothymol blue was not considered to be suitable for our purposes. Instead, we developed two assays suitable for 384-well format using phosphatidylcholine as a substrate, with either cresol red (CR) as a color pH indicator or fluorescein as a fluorescent pH indicator.

### 3.1 PLA_2_ activity assay based on cresol red

#### 3.1.1 Assay optimization

Going from pH 8.8 to 7.0, protonation of the pH indicator cresol red results in a color change from red to yellow. When cresol red is added to the assay reagent mixture, PLA_2_ activity can be detected as a decrease in absorbance at 572 nm.[25, 27] The pH of the assay solution was first adjusted to 8.0 in order to allow sensitive detection of the pH shift. Optimal conditions were studied and achieved in several optimization steps, as described below. More detailed information and results on the development of the cresol red-based PLA_2_ activity assay are provided in the supporting information S2 and S3.

The optimal concentrations for cresol red and the substrate phosphatidylcholine were studied by performing serial dilution experiments. *Daboia russelii russelii* (DRR) snake venom, which is known for its relatively high content of svPLA_2_, was used as a test venom for assay optimization at a concentration of 12.5 µg/mL.[11] Figure 1 shows time-course absorbance measurements in a cuvette using optimized assay conditions and DRR venom.

**Figure 1:**
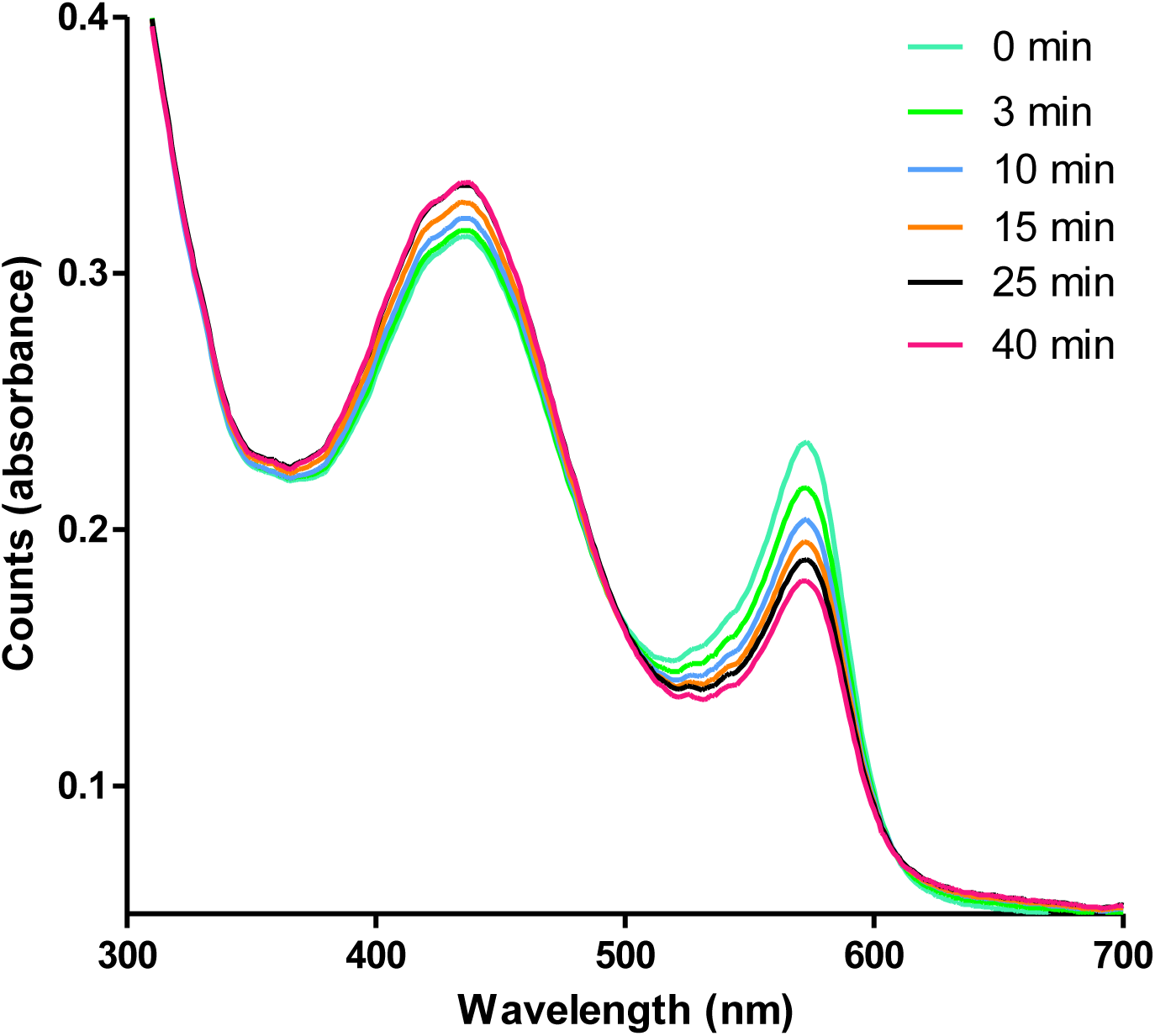
Absorbance spectra obtained during progression of the cresol red-based PLA_2_ activity assay. Conditions: phosphatidylcholine concentration, 0.66 mM; cresol red concentration, 37 µM; DRR venom concentration, 12.5 µg/mL.

The color change of cresol red over time is observed as an increasing absorbance at 430 nm and a decreasing absorbance at 570 nm. As the absolute change is most substantial at 570 nm, this wavelength was used for the readout of the cresol red-based PLA_2_ assay. Higher concentrations of cresol red resulted in more intense red coloring of the assay mixture, detected as a higher absorbance at 572 nm at the start of the measurement (Figure S2). The optimal cresol red concentration was found to be 37 µM. Increasing the concentration of phosphatidylcholine resulted in an increased acidification of the assay mixture upon assay progression when svPLA_2_s were present. The optimal substrate concentration was determined to be 0.66 mM. Concentrations of 0.66 mM and 37 µM for phosphatidylcholine and cresol red, respectively, were used in all subsequent experiments.

#### 3.1.2 Assay evaluation

The optimized assay was evaluated for sensitivity and limit of detection (LOD) by analyzing different concentrations of DRR snake venom.

The concentration-response plot for svPLA_2_ activity was assessed by analyzing a serial dilution series of DRR venom in water (freeze-dried after addition to the wells) with final assay concentrations of 50, 25, 12.5, 6.3 and 3.2 μg/mL (Figure 2).

**Figure 2.**
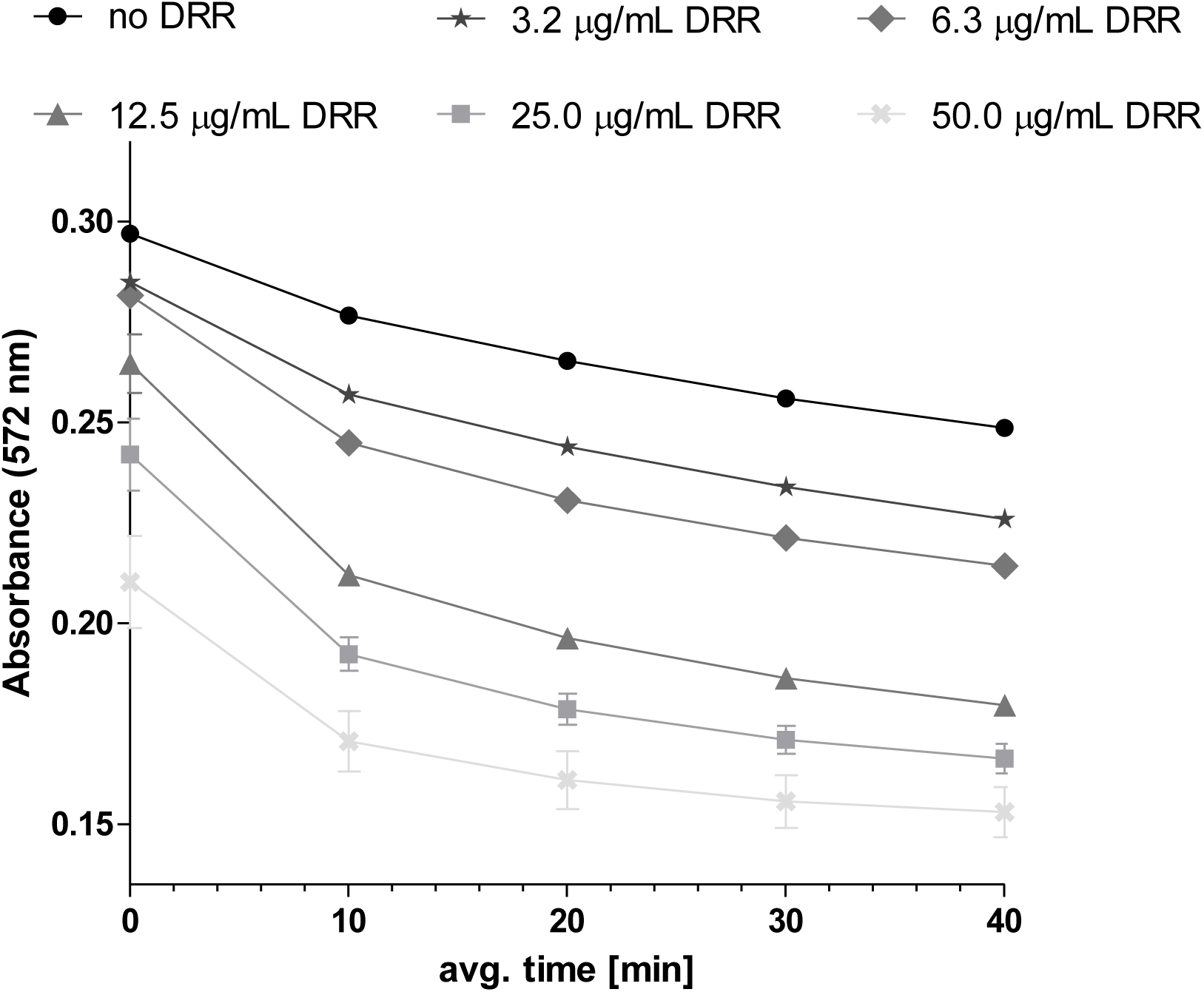
Kinetic absorbance measurements obtained of the cresol red based PLA_2_ assay. Concentrations of phosphatidylcholine and cresol red were 0.66 mM and 37 μM, respectively. Assay was performed in presence of a DRR concentration series with final assay concentrations of 50 μg/mL, 25 μg/mL, 12.5 μg/mL, 6.3 μg/mL and 3.2 μg/mL (visualized from bottom to top). The absorbance was set at 570 nm. Each curve represents the mean of two measurements and the error bars represent SEMs

The starting steepness of the declining curve is correlated to the DRR venom concentration and thus the overall activity of svPLA_2_s present in the assay mixture. Note that the starting point of the kinetic measurement is the moment the plate reader measurement is initiated and thus does not represent the real starting point, which is the moment the assay mix is pipetted to the venom. As can be seen, increasing DRR concentrations resulted in faster declines in absorbance over time (Figure 2). Also, the higher the concentration of DRR the lower the absorbance of the first measured point due to the initial faster enzymatic conversion rates. Therefore, after pipetting the assay mix to a well plate, the assay readout was started directly.

The assay specificity, i.e. determining whether the assay reflects the action of PLA_2_s as the cause of the detected acidification, was performed by analyzing DRR venom in presence of the PLA_2_ inhibitor varespladib. A recent study by Lewin *et al*. demonstrated methyl-varespladib and varespladib to reduce venom PLA_2_-induced *in vivo* pathologies.[17] These compounds were shown to be potent inhibitors for a multitude of svPLA_2_s *in vitro.* An IC50 value of 0.96 µM ± 0.04 µM was determined for varespladib.[17] As varespladib is a broad-range (non-specific) PLA_2_ inhibitor, it was anticipated to be able to inhibit the majority of svPLA_2_s present in a venom and as such concentration-dependently reduce the hydrolyzation rate of phosphatidylcholine in the assay. The effect of the concentration of varespladib on the activity of DRR (final assay concentration, 12.5 µg/mL) was determined using a serial dilution series of varespladib in Tris buffer (1 mM; pH 8; 10 µl/well) with final concentrations of 20, 2, 0.002, and 0 μM (Figure 3).

**Figure 3:**
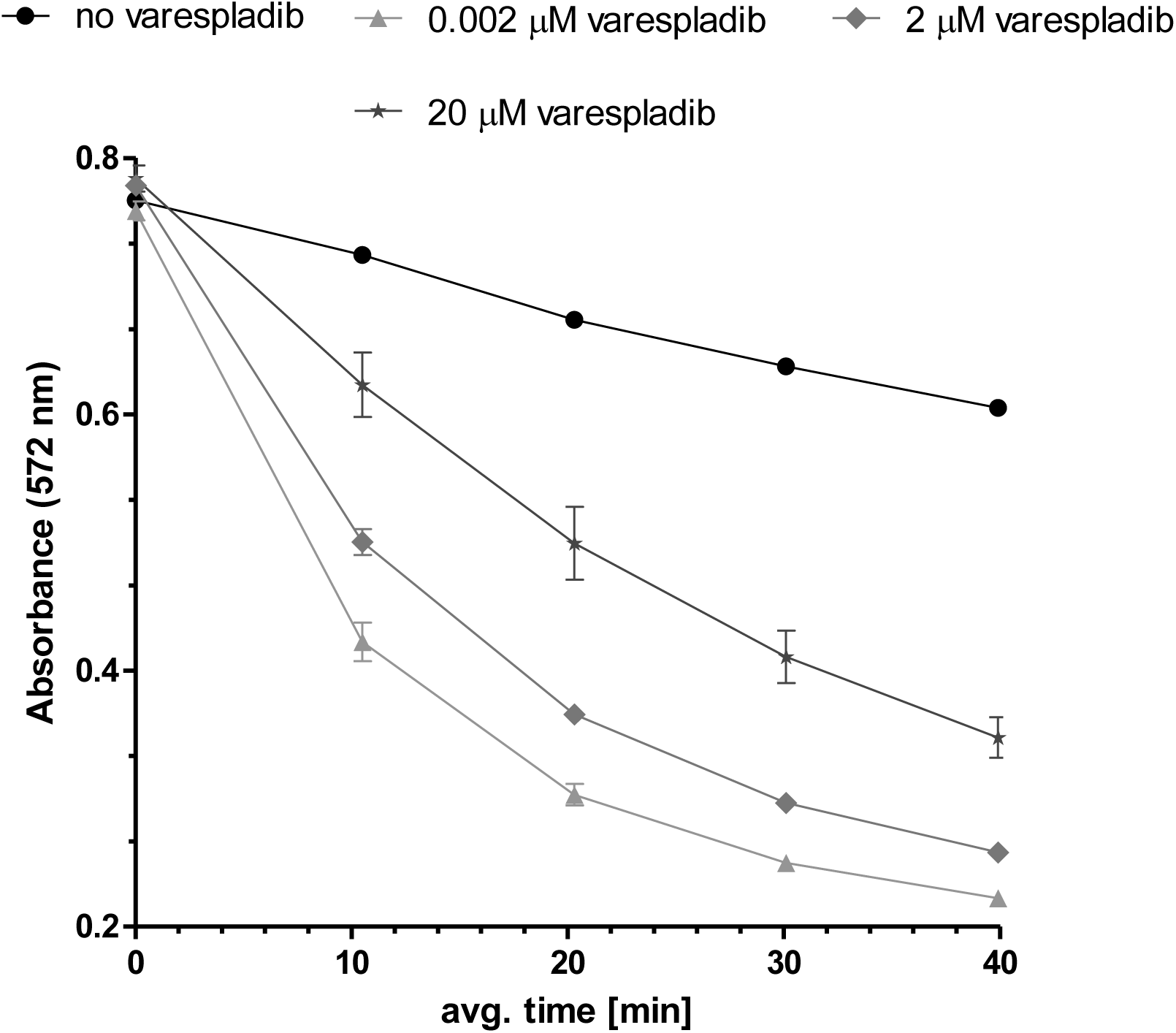
Kinetic absorbance measurements obtained for assessing the inhibitory effect of the PLA_2_ inhibitor varespladib on the cresol red based PLA_2_ assay. Concentrations of phosphatidylcholine, cresol red, and DRR were 0.66 mM and 37 μM, and 12.5 μg/mL, respectively. Assay was performed in presence of varespladib with final concentrations of 20, 2, 0.002, and 0 μM. The absorbance was set at 570 nm. Each curve represents the mean of three measurements and the error bars represent SEMs.

When increasing concentrations of varespladib were added to the assay mixture, a concentration-dependent decrease in PLA_2_ activity was observed visible as a decrease in steepness of the curves. The highest concentration varespladib tested almost overlapped with the blank measurement (upper curve) clearly showing that the enzymatic acidification of the bioassay mixture was almost fully inhibited at this concentration varespladib. In the blank measurement, a decrease in absorbance was observed due to non-enzymatic (chemical) hydrolysis of phosphatidylcholine.

### 3.2 PLA_2_ assay based on fluorescein

#### 3.2.1 Assay development

The fluorescence quantum yield of fluorescein is pH dependent, being highest at around pH 8. Its fluorescence decreases in the presence of lowered pH, which is the detection principle of the fluorescence assay format. Below pH 6 the emission starts to get weak, and becomes undetectable at very acidic conditions.[30, 31] For this assay a Tris buffer of pH 8 was chosen (as for cresol red) and the decrease of fluorescence intensity of fluorescein was monitored as result of PLA_2_ activity causing a drop in pH.[30, 31] Additional information and results on the optimization of the fluorescein-based PLA_2_ activity assay is provided in Section S4 of the supporting information. Different fluorescein concentrations were tested (final concentrations: 0.2, 0.5, 1.0, and 5.0 µM). The same phosphatidylcholine concentration as for the cresol red-based assay was used assuming the substrate concentration to be independent of the pH indicator used. Increasing the fluorescein concentration yielded more intense fluorescence signals at the start of a measurement (see Figure S4 in the supporting information). The optimal fluorescein concentration was determined to be 1 µM, providing optimal signal-to-noise ratios while achieving acceptable assay repeatability. Concentrations of 1 µM fluorescein and 0.66 mM phosphatidylcholine were used for all subsequent experiments.

#### 3.2.2 Fluorescein PLA_2_ assay evaluation

The optimized fluorescein-based assay was further evaluated for sensitivity and LOD. DRR concentrations of 12.5, 6.1, 3.0, 1.6, 0.8, and 0 µg/mL in Tris buffer (1 mM; pH 8; 10 µl/well) were analyzed (Figure 4). The specificity was again tested by performing the activity assay with DRR in presence of different concentrations of varespladib.

**Figure 4:**
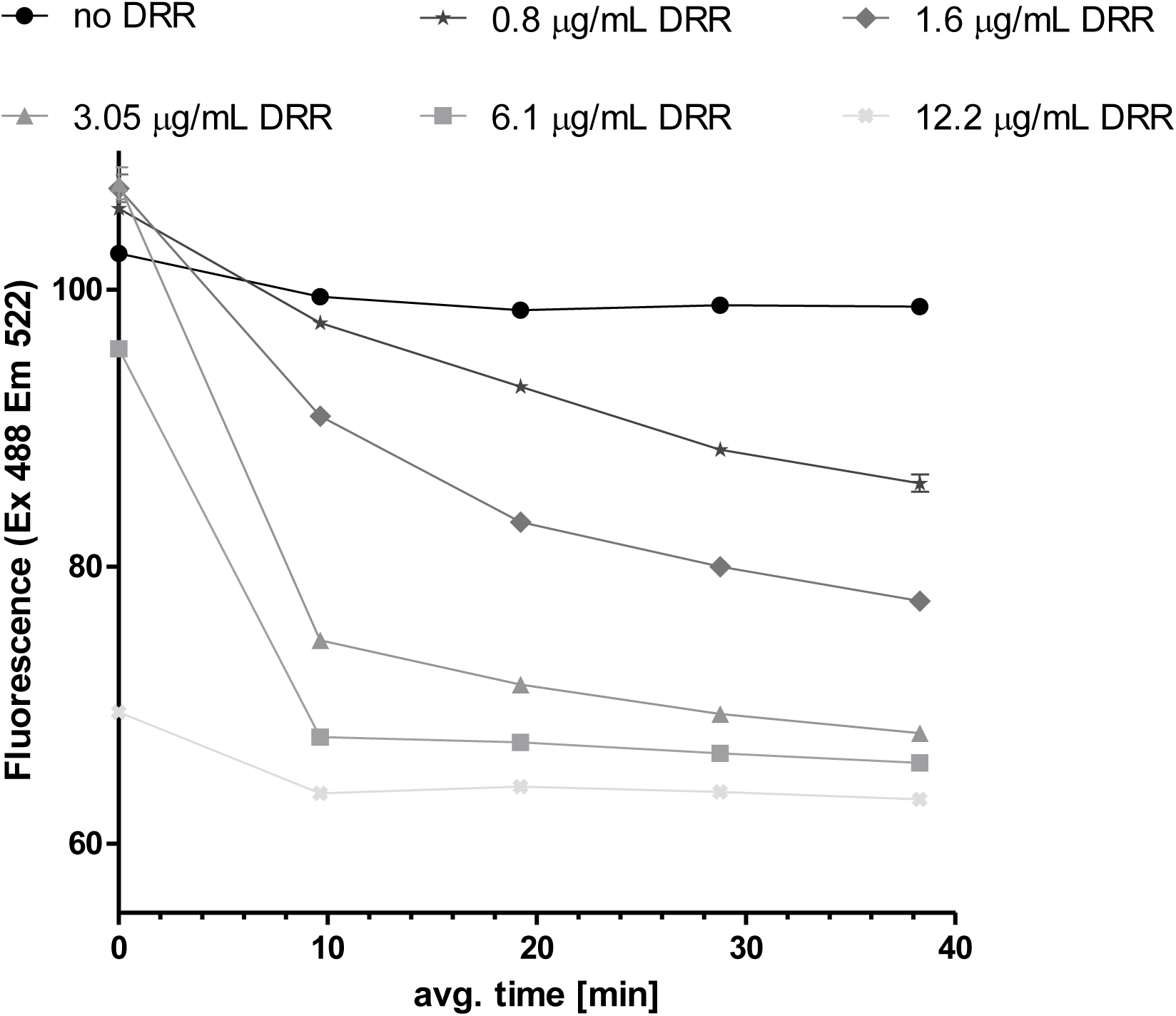
Kinetic fluorescence measurements obtained for the fluorescein based PLA2 assay. Concentrations of phosphatidylcholine and fluorescein were 0.66 mM and 1 μM, respectively. DRR concentrations tested: 12.5, 6.3, 3.2, 1.6, and 0.8 μg/mL. Fluorescence was measured using an excitation wavelength of 488 nm and an emission wavelength of 520 nm. Each curve represents the mean of three measurements and the error bars represent SEMs.

The DRR serial dilution series was added to different wells and then vacuum centrifuge dried prior to assay pipetting to avoid undesired dilution effects. As with the previous assay, the starting point of the kinetic measurement is the moment the plate reader measurement is initiated and as such thus not the real starting point, which is the moment the assay mix is pipetted to the venom. Therefore, after pipetting the assay mix to a well plate, the assay readout was started as soon as possible. Increased DRR concentrations resulted in increased progression of the svPLA_2_s enzymatic activity visible as declined fluorescent intensities observed at the first measurement point of the curves. The last measurement point for the two highest DRR concentrations tested (12.5 µg/mL and 6.3 µg/mL) almost overlapped, indicating a smaller assay window for the fluorescein-based assay as compared to the cresol red assay when performing an end-point measurement. The results also show that the fluorescein assay was more sensitive than the cresol red assay for DRR venom. With this assay, clear hydrolyzation was already observed within 10 minutes at a DRR concentration of 1.6 μg/mL. In the fluorescein assay, a concentration of 0.8 μg/mL DRR still gave a measurable effect after 40 minutes.

The fluorescein-based assay was then evaluated for specificity for PLA_2_ activity by measuring a concentration series of the PLA_2_ inhibitor varespladib. The effect of the concentration of varespladib on the activity of DRR was determined for two final concentrations of DRR snake venom (12.5 and 1.6 µg/mL; Figure 5 & 6 respectively). The choice to test two DRR concentrations was because 12.5 µg/mL (same concentration as used for the cresol red assay) gave very rapid enzymatic hydrolyzation for this assay. Serial dilution series of varespladib in Tris buffer (1 mM; pH 8; 10 µl/well) with final concentrations of 50, 5.0, 0.5, 0.05, and 0 μM were tested (Figures 5 and 6). Varespladib was found to inhibit the PLA_2_ activity in a concentration-dependent manner, with full inhibition observed at a concentration of 5 µM for the 12.5 µg/mL DRR experiment and 50 µM for the 1.6 µg/mL DRR experiment. As also observed with the cresol red assay, the fluorescein-based assay thus shows specificity towards enzymatic PLA_2_ activity.

**Figure 5:**
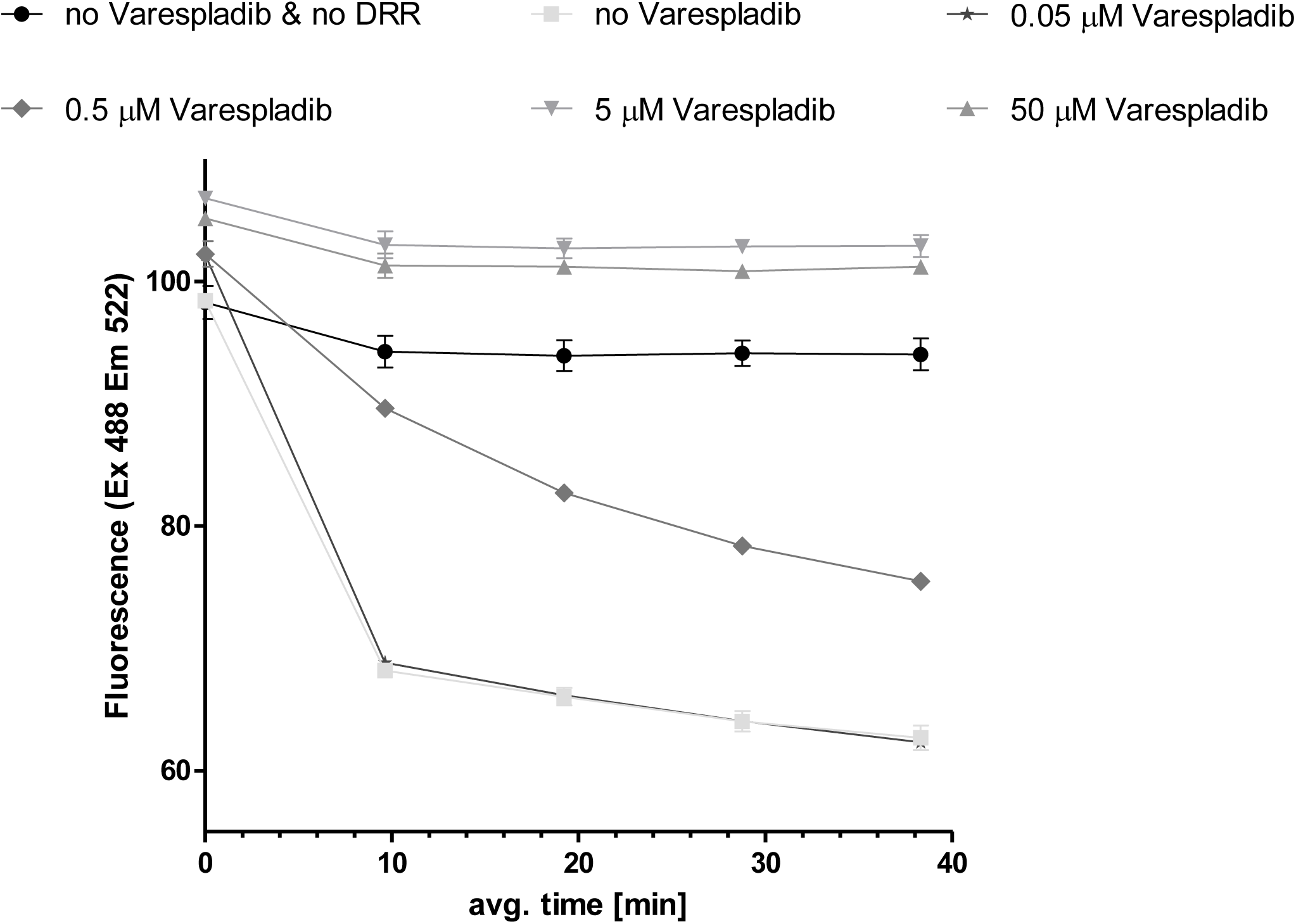
Kinetic fluorescence measurements obtained for assessing the inhibitory effect of the PLA_2_ inhibitor varespladib on the fluorescein based PLA_2_ assay. Concentrations of phosphatidylcholine, fluorescein, and DRR were 0.66 mM and 1 μM, and 12.5 μg/mL, respectively. The assay was performed in presence of varespladib with final concentrations of 50, 5.0, 0.5, 0.05, and 0 μM. Fluorescence was measured using an excitation wavelength of 488 nm and an emission wavelength of 520 nm. Each curve represents the mean of two measurements and the error bars represent SEMs.

**Figure 6:**
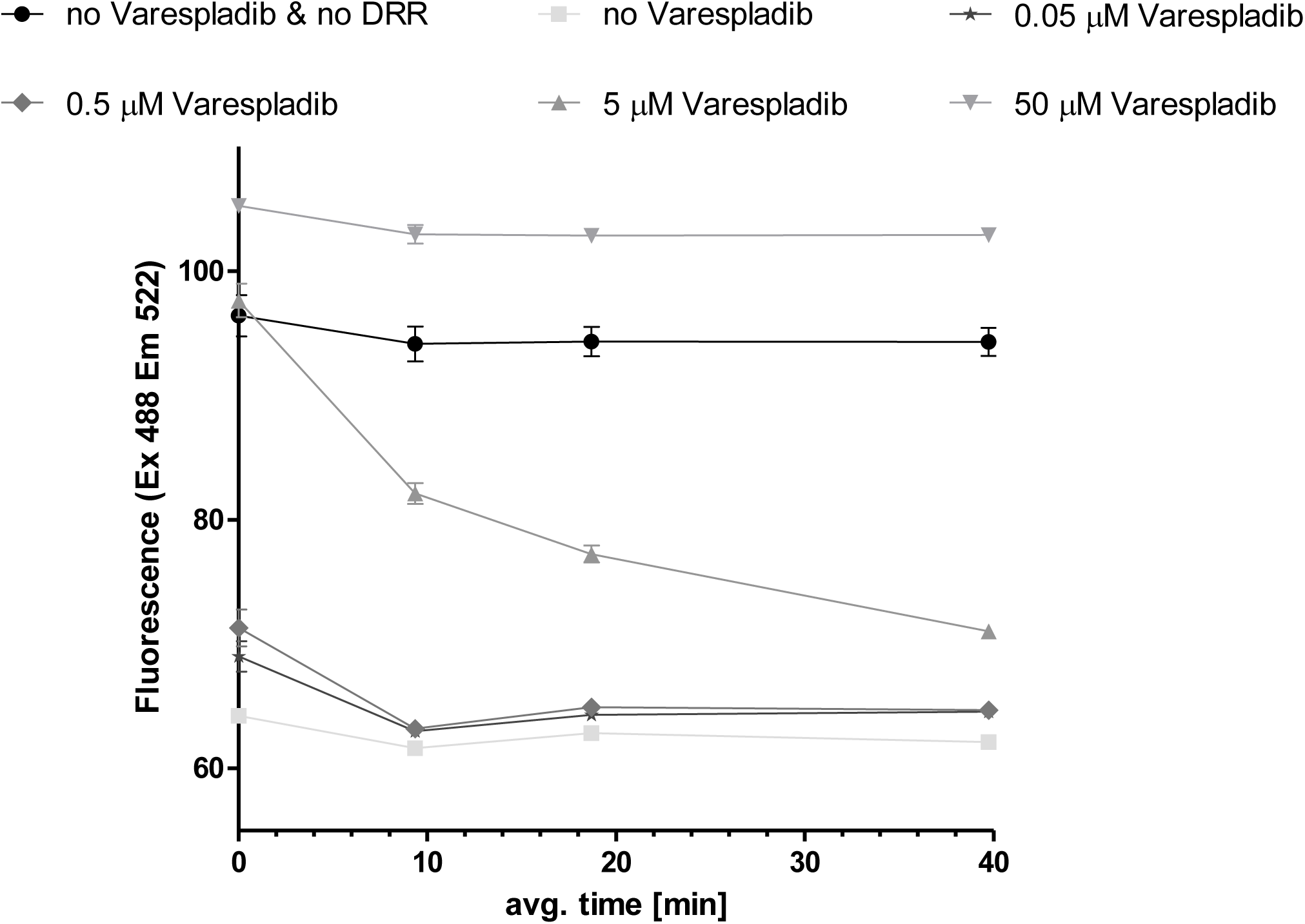
Kinetic fluorescence measurements obtained for assessing the inhibitory effect of the PLA_2_ inhibitor varespladib on the fluorescein based PLA_2_ assay. Concentrations of phosphatidylcholine, fluorescein, and DRR were 0.66 mM and 1 μM, and 1.6 μg/mL, respectively. The assay was performed in presence of varespladib with final concentrations of 50, 5.0, 0.5, 0.05, and 0 μM. Fluorescence was measured using an excitation wavelength of 488 nm and an emission wavelength of 520 nm. Each curve represents the mean of two measurements and the error bars represent SEMs.

To summarize the two approaches compared here, the PLA_2_ assay using cresol red facilitates measuring a wider range of PLA_2_ concentrations in comparison with the fluorescein-based assay. However, the latter assay is more sensitive.

### 3.3 Coupling of the two PLA_2_ assay formats with nanofractionation analytics

Finally, the two developed assays were incorporated into the nanofractionation analytics platform. This allowed for obtaining bioactive PLA_2_ profiles of snake venoms after chromatographic separation.

svPLA_2_s are derived from members of the sPLA_2_ family, which are found in mammals and other vertebrates. It is said that the genes expressing svPLA_2_s have undergone gene duplication and accelerated evolution over evolutionary time, which has resulted in structural, functional, and expressional level variations of svPLA_2_s isoforms.[14, 32] As such, many snake venoms are known to contain a number of different svPLA_2_ isoforms. The presence of these isoforms, and their abundances, can thus vary greatly according to species, but also according to, for example, population, age and sex of a snake, and (prey) ecology.[33–35] The generic svPLA_2_ tertiary structure, may contain up to seven disulfide bridges. This means that for an enzyme, svPLA_2_s are considered to be rather stable and are therefore relatively resistant to heat, organic solvents, and acidic conditions.[17] These properties, their high abundance in many snake venoms, and their relatively low molecular weight (∼13 to 15 kDa) is advantageous for their intact isolation/fractionation using reversed phase liquid chromatography (RPLC) thereby allowing subsequent characterization and bioactivity profiling.[36, 37]

The nanofractionation analytics platform connects chromatographic separations of venoms with UV data collection, optionally mass spectrometry, and high-resolution fractionation on well plates followed by bioassaying. This platform for screening snake venoms has been described earlier by others, including for example Mladic et al.[29] and Still et al.[28] for other assay formats. In this study, snake venoms were separated using RP-LC with a post column split directing 90% of the effluent to a high-resolution fraction collector (e.g. the FractioMate) and 10% to UV and optionally MS. After fraction collection in 384-well plates, the plates were vacuum centrifuged to remove solvents after which one of the two PLA_2_ bioassays was performed. After bioassaying, reconstructed bioassay chromatograms were plotted which could then be correlated to UV and MS traces in an effort to identify the enzymatically active svPLA_2_s.

Venom of six snake species were screened for svPLA_2_ activity using the cresol red PLA_2_ assay and venoms of two snakes were screened using the fluorescein-based PLA_2_ assay. The results obtained for the cresol red based assay are shown in Figure 7. Each experiment was performed in duplicate and both the LC-UV spectral data as well as the chromatographic bioactivity profiles (representing the mean of three measurements) are provided. The bioassay chromatograms were constructed by plotting the calculated slopes of each measured svPLA_2_ activity curve of each well against fractionation time. As 6-sec fractions were collected, high resolution reconstructed bioassay chromatograms were obtained. svPLA_2_ activity is detected as an increase in the slopes of the kinetic measurements and is svPLA_2_ concentration-dependent (e.g. as shown in Figure 3). The enzymatic activities of eluted svPLA_2_s are therefore visible in the reconstructed bioassay chromatograms as positive peaks. Each venom analyzed was found to have its own characteristic fingerprint or signature-like LC-UV chromatogram with corresponding bioactivity chromatogram profile. The results displayed in Figure 7 show that the bioactive peaks for all venoms analyzed eluted in the 15 to 25-minute time frame. Later eluting (non-bioactive) peaks may, although not likely for the relatively stable svPLA_2_s, be denaturized svPLA_2_s (due to the high organic solvent (i.e. acetonitrile) concentration during elution). As can be seen from the chromatographic PLA_2_ bioactivity data, all the snake venoms analyzed show positive svPLA_2_ bioactivity peaks, which indicates that all venoms profiled contain at least one, but in most cases multiple, svPLA_2_s.

**Figure 7:**
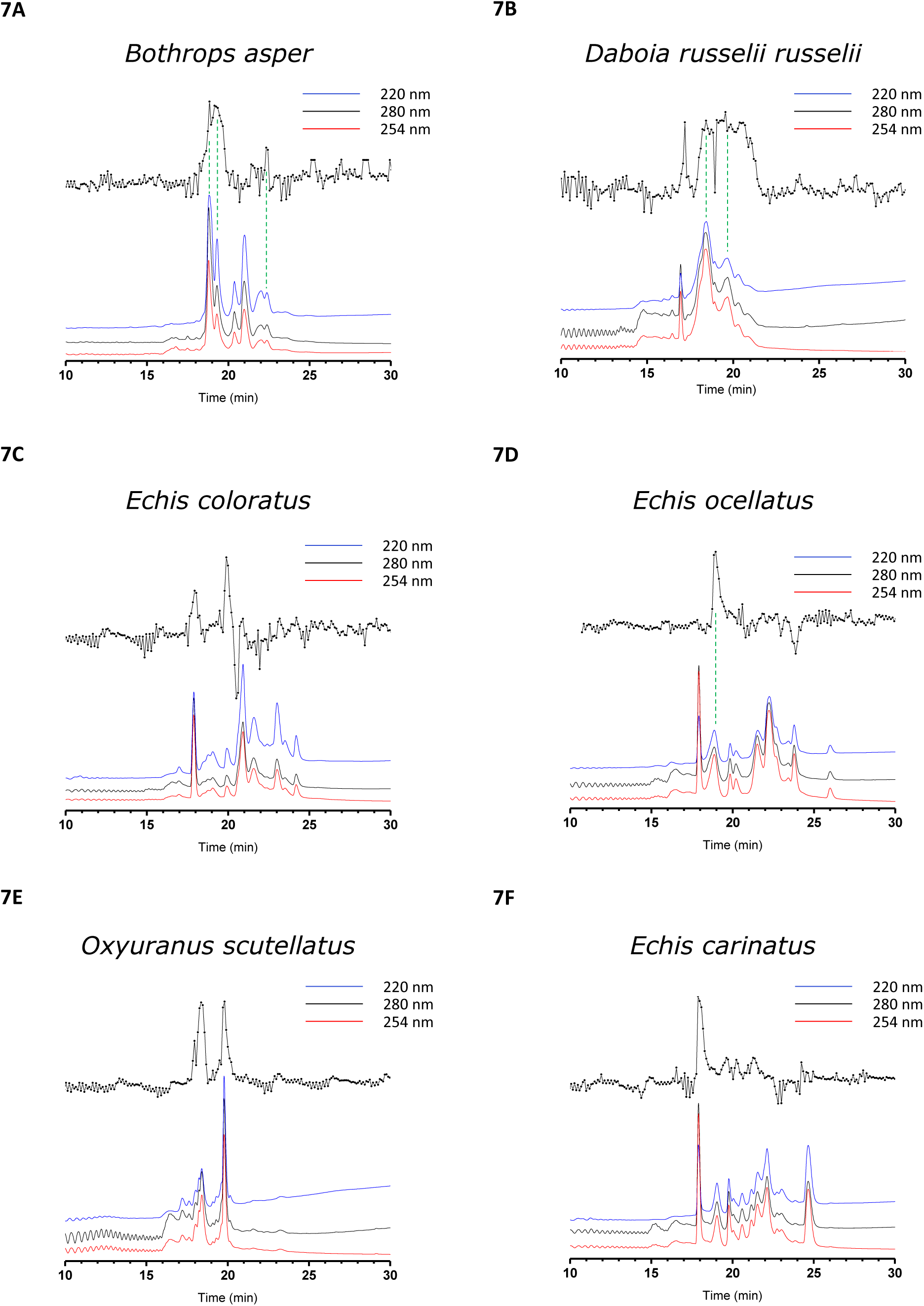
UV-chromatograms measured at three different wavelengths (lower three superimposed chromatograms per figure) and corresponding cresol red assay-PLA_2_ reconstructed bioassay chromatograms (upper superimposed chromatograms per figure) obtained from RP-LC separated snake venoms. Snake venoms measured: (a) *Bothrops asper;* (b) *Daboia russelii russelii; (c) Echis coloratus; (d) Echis ocellatus; (e) Oxyuranus scutellatus; (f) Echis carinatus*. Experimental conditions: Snake venom concentrations injected were 1 mg/mL with an injection volume of 50 uL. A UV-DAD detector measured the spectrum between 200 and 300 nm of which 220 nm (blue), 280 nm (black) and 254 nm (red) are plotted in the figures. For correlating bioactivity peaks to tentative toxin identification, the reader is referred to section 3.4 (only for the venoms of *Bothrops asper, Daboia russelii russelii*, and *Echis ocellatus* this tentative toxin identification was performed). Green dotted lines indicate tentatively identified svPLA_2_s.

With the fluorescein-based PLA_2_ assay, *Bothrops asper* was profiled (Figure 8). For the fluorescein-based assay it was observed that larger assay windows in combination with lower fluctuation in the baselines were obtained as compared to the cresol red based assay, visualized as positive bioactivity peaks in the fluorescence bioassay chromatograms. For *Bothrops asper* venom, several svPLA_2_ bioactivities are clearly observed and, when compared to the results obtained from the cresol red assay, additional bioactivities were visualized due to the higher sensitivity of the fluorescein-based assay. After the first set of closely co-eluting svPLA_2_s observed in both assays (between ∼17 to 19 minutes), these additional svPLA_2_ activities are observed after the first broad peak and eluted as one sharp peak followed by two non-baseline separated bioactivity peaks.

**Figure 8:**
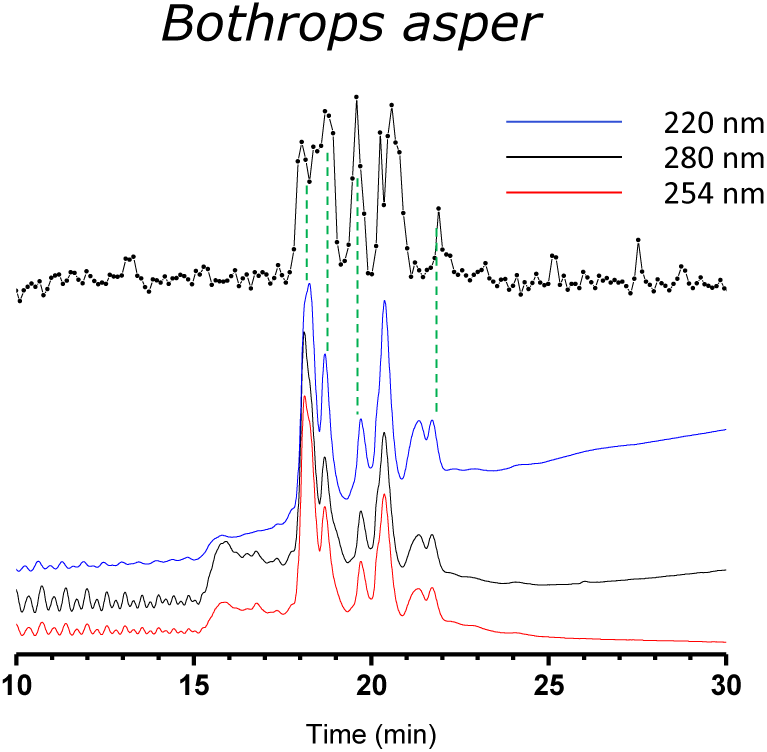
UV-chromatograms measured at three different wavelengths (lower three superimposed chromatograms per figure) and corresponding fluorescein assay-PLA_2_ reconstructed bioassay chromatograms (upper superimposed chromatograms per figure) obtained from RP-LC separated snake venoms. Snake venom measured: *Bothrops asper;* Experimental conditions: Snake venom concentrations injected were 1 mg/mL with an injection volume of 50 µL. A UV-DAD detector measured the spectrum between 200 and 300 nm of which 220 nm (blue), 280 nm (black) and 254 nm (red) are plotted in the figures. For correlating bioactivity peaks to tentative toxin identification, the reader is referred to section 3.4. Green dotted lines indicate tentatively identified svPLA_2_s.

It has to be noted that for both assays, not all positive peaks observed need to correspond with svPLA_2_ activities. Although not expected, other toxins in venoms could have enzymatic activities directly or indirectly resulting in acidification or basification of the assay medium. As the assay mixture’s color change is dependent on the pH indicator and therefore the pH of the mixture, compounds with acidic and/or basic functional groups present in high concentrations could potentially chemically modulate the assay pH and as such result in negative or positive peaks. Although in general this is not likely to occur, this might be the outcome of the negative peak observed in the bioactivity chromatogram of *Echis coloratus* (Figure 7C). This negative activity might have resulted from a compound having a basifying effect on the assay mixture’s pH. In this specific case it is not expected to have resulted from an activity linked to enzymatic PLA_2_ activity. In addition, it is known that svPLA_2_s can display a change in activity (mostly an increase) when enzymatically converting higher-ordered lipid aggregate substrates as compared to monomeric substrates.[38] As such, some svPLA_2_s may show low activity for the phospholipid substrate used in this study.

### 3.4 Correlation of bioactive peaks with MS and proteomics data for toxin identification

From literature and the database Uniprot (https://www.uniprot.org/)[39] it was found that all snake venoms profiled in this study have multiple known svPLA_2_s. Table 2 displays a summarized list of the number of svPLA_2_s found in the literature and from the Uniprot database. The table also lists the number of PLA_2_-linked enzymatic activities found for both assays developed and applied in this study after nanofractionation of the profiled venoms. From the overview it is evident that for most snake venoms analyzed, more individual svPLA_2_ isoforms are reported in the literature and Uniprot, when compared to the number of individual positive peaks found per venom in this study. The reason for this is likely as follows: firstly, snake venoms of the same species, but from different geographical origins, often have different venom compositions implying that not all svPLA_2_s necessarily will be expressed at significant levels in each venom. Secondly, the chromatographic overlap of svPLA_2_ isoforms resulting from chromatographic co-elution of (especially structurally similar) svPLA_2_s results in overlapping bioactivity peaks for which multiple svPLA_2_ isoforms can be assigned to. This was also clearly shown in other studies by Still et al.[19] and Slagboom et al.[40] In addition, some svPLA_2_s in Uniprot could be present in a given venom but showed reduced catalytic activity or have lost their activity due to denaturation in our experiments. Furthermore, some sub-classes of svPLA_2_s have lost their enzymatic activity (among which are neurotoxic svPLA_2_s and lysine/serine 49 svPLA_2_s) and/or only have minor enzymatic activity as compared to their other biological functions which can include specific target-binding, chaperoning, and/or membrane-disrupting functions.[13]

**Table 2:** Overview of numbers of svPLA_2_ isoforms identified by different research groups and placed in the Uniprot database, numbers of svPLA_2_ isoforms identified by Slagboom *et al*, and detected number of bioactives in the developed cresol red and/or fluorescein-based PLA*_2_* assays described in this study. n.d. means no data available.

| Snake species | Uniprot database svPLA<sub>2</sub> isoforms | Masses of svPLA<sub>2</sub>s identified by Slagboom *et al.* | svPLA<sub>2</sub> bioactivities found in cresol red PLA<sub>2</sub> assay | svPLA<sub>2</sub> bioactivities found in fluorescein PLA<sub>2</sub> assay |
| --- | --- | --- | --- | --- |
| *Bothrops asper* | 6 | 4 | 2 | 6 |
| *Daboia russelii russelii* | ~33 | 2 | n > 3 | n.d. |
| *Echis coloratus* | 10 | n.d. | 2-3 | n.d. |
| *Echis ocellatus* | 1 | 2 | 1 | n.d. |
| *Oxyuranus scutellatus* | 12 | 0 | 2-5 | n.d. |
| *Echis carinatus* | 6 | n.d. | 1 | n.d. |

Accurate mass measurements and proteomics data of the toxins present in several snake venoms included in this study were recently reported by Slagboom *et al.*[40] They used the same LC-nanofractionation platform in combination with parallel acquired MS data and profiled coagulopathic activity followed by proteomic characterization of the active coagulopathic compounds. The following three snake venoms analyzed by Slagboom *et al* were also investigated in this study: Bothrops asper, Echis Ocellatus, and Daboia russelii russelii. The MS and proteomics data of Slagboom *et al.* was used here for correlation of svPLA_2_ bioactivity to hypothetical toxin identification in order to determine platform applicability for svPLA_2_ profiling of snake venoms towards identification of the bioactives. Slagboom *et al.* profiled and tentatively identified venom toxins that exhibited coagulopathic activity and thus might not match svPLA_2_ activities measured this study. Several positive signals in our svPLA_2_ assays could, however, be correlated to identified svPLA_2_s by Slagboom *et al.,* as they fell within the same retention timeframes (note that the same chromatographic conditions were used in both studies). In *Bothrops asper*, Slagboom *et al.* found four svPLA_2_s with *m/z*-values of 1378.3697^10+^, 1373.3688^10+^, 1267.7906^11+^, and 1164.8881^12+^. Combining the fluorescein and cresol red PLA_2_ assay data of *Bothrops asper* measured this study after nanofractionation resulted in correlation of most activities with the MS and proteomics data from Slagboom *et al*. The first two eluting svPLA_2_s with *m/z*-values of 1378.3697^10+^ and 1373.3688^10+^co-eluted in the chromatogram and therefore could not be differentiated. Two additional activities were detected in the fluorescent PLA_2_ assay, which were not measured by Slagboom *et al (*who only did MS and proteomics analysis on coagulopathic venom toxins). Two masses within the svPLA_2_ mass range were identified by Slagboom *et al* for *Daboia russelii russelii* venom and one for *Echis ocellatus* venom. For both venoms, the identified *m/z*-values could be correlated to the svPLA_2_ bioactivity peaks falling within the same retention timeframe and were 1511.6962^9+^ and 1518.5946^9+^ for *Daboia russelii russelii* venom, and 1537.0489^9+^ for *Echis ocellatus* venom. In Figure 7 and Figure 8, the tentatively identified svPLA_2_s correlating to bioactivity peaks are indicated with green marker arrows pointing to their respective bioactivity peaks for *Bothrops asper*, *Echis ocellatus*, and *Daboia russelii russelii* venoms.

## 4. Conclusion

The PLA_2_ assays developed herein were demonstrated to be sensitive and robust and were successfully coupled to nanofractionation analytics to analyze svPLA_2_ activity of separated venom toxins. The PLA_2_ assay using cresol red was found to have the largest assay window, i.e. being able to detect a larger variation in PLA_2_ concentrations, while the fluorescein-based assay proved to be the most sensitive assay, thereby making both assays complementary to one another. When using C18 RP-LC with nanofractionation analytics for svPLA_2_ profiling of snake venoms, reproducible specific fingerprint-like separation profiles were obtained of which bioactive svPLA_2_s showed positive bioactivity peaks in the reconstructed bioactivity chromatograms. As many svPLA_2_s have similar primary sequences, and thus much structural resemblance, overlap of different bioactive svPLA_2_s by co-elution was expected and was indeed observed in this study. Prior accurate mass and proteomics data of some of the venoms used in this study was repurposed here for svPLA_2_ identification, and to assess assay and analytical platform performance, resulting in bioactivity chromatograms with tentatively identified enzymatically active svPLA_2_s. We anticipate that our development of new analytics for rapidly profiling svPLA_2_ activity will have utility for future research on snakebite pathologies caused by svPLA_2_ toxins, and for the identification of novel inhibitory molecules capable of neutralizing svPLA_2_s for their future selection and translation into snakebite therapeutics.

## Acknowledgements

This study was supported by: (i) a Sir Henry Dale Fellowship to N.R.C.736 (200517/Z/16/Z) jointly funded by the Wellcome Trust and Royal Society, and (ii) a China Scholarship Council (CSC) fellowship to C.X.

