## Supplemental file for "Development of high-throughput screening assays for profiling snake venom Phospholipase A_2_ activity after high-resolution chromatographic fractionation"

***S1. Typical kinetic curve results obtained when running the PLA2 assays***

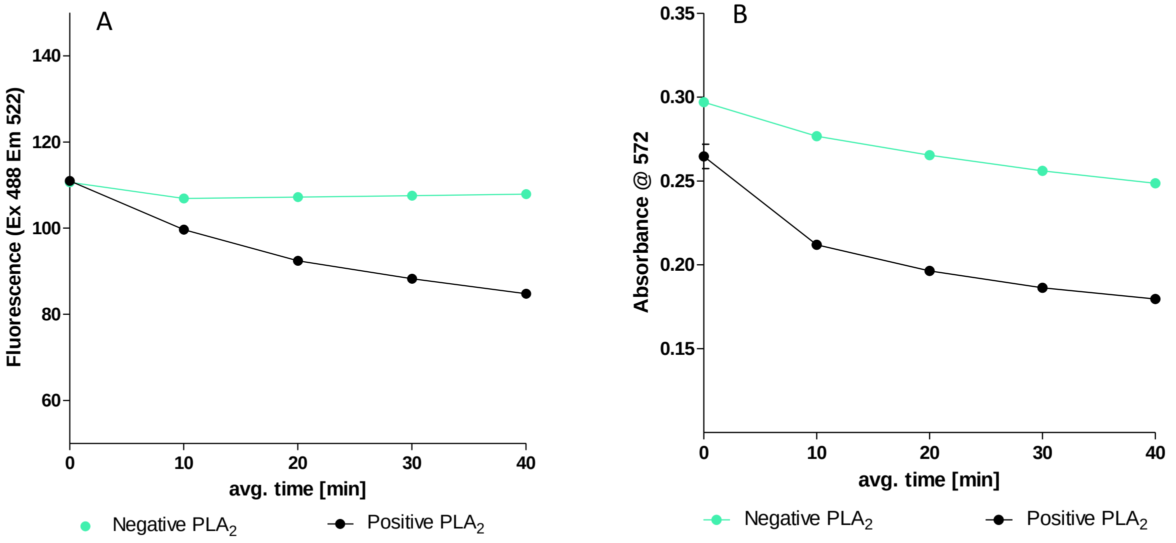


Figure S1. Typical results of running both $PLA_2$ assay formats with and without a venom rich in $PLA_2$s (i.e. DRR venom). a) The assay using fluorescein, and b) the assay using cresol red. In both figure S1A and S1B, the green curve is the average blank measurement (i.e. no snake venom) and the black curve is the average $PLA_2$ activity measurement (i.e. with snake venom). As can be seen from both assay results, the blank curves also show a slight decrease in absorbance or fluorescence, which is due to hydrolysis of phosphatidylcholine upon contact with water.

***S2. Serial dilution of cresol red***

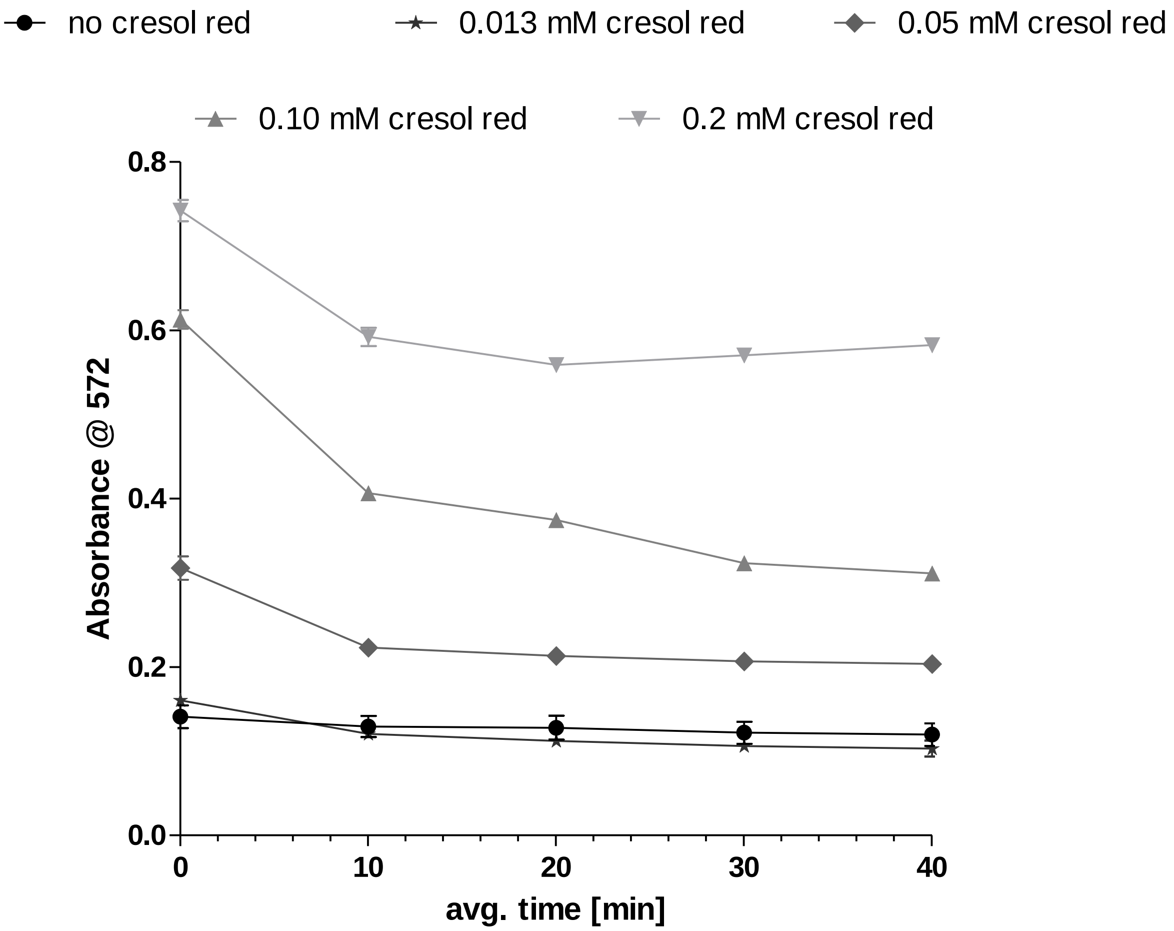


Figure S2. Optimization of the cresol red assay concentration in the cresol red based $PLA_2$ assay. The different assay concentrations tested were 0.2, 0.10, 0.05, 0.013, and 0.0 mM cresol red final concentration in presence 12.5 µg/mL DRR venom, respectively.  Each curve represents the mean of three measurements and the error bars represent SEMs. Increasing the cresol red concentrations gives an increased absorbance signal. The optimal final cresol red concentration was found to be 0.05 mM, due to the minimal decrease in combination of steepness of the curve, absorbance level and most reproducible results. In these tests, the final concentration of phosphatidylcholine and DRR was kept constant at 0.66 mM and 12.5 µg/mL, respectively.

***S3. Serial dilution of phosphatidylcholine***

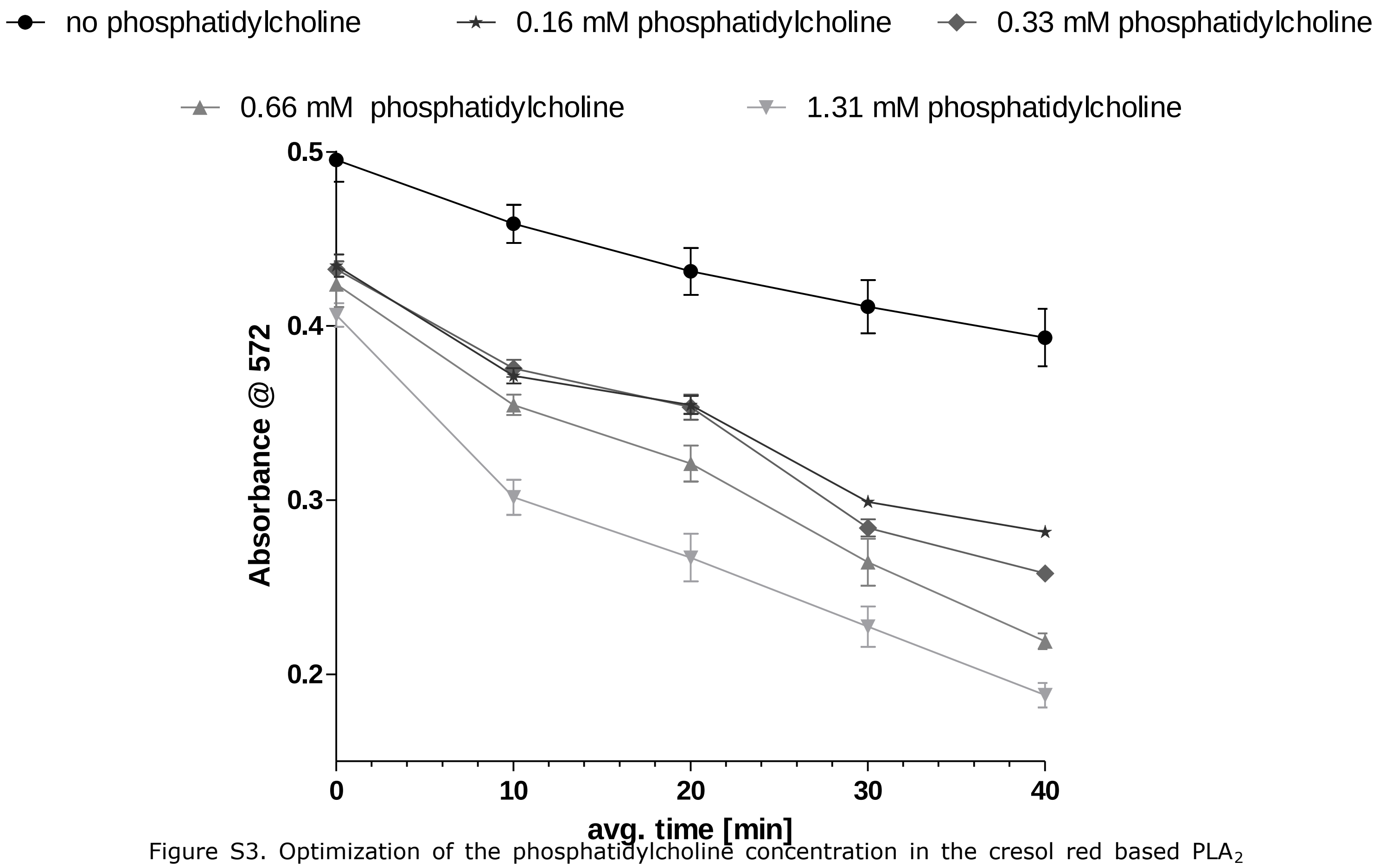


Figure S3. Optimization of the phosphatidylcholine concentration in the cresol red based $PLA_2$ assay. The different assay concentrations tested were 0.0, 0.16, 0.33, 0.66, and 1.13 mM phosphatidylcholine in presence of 0.05 mM and 12.5 µg/mL cresol red and DRR venom, respectively. In the figure, each curve represents the mean of three measurements and the error bars represent SEMs. Increasing the phosphatidylcholine concentration gives an increased decline of the absorbance signal, which can be correlated with the increased velocity of acidification of the assay medium. The optimal final phosphatidylcholine concentration was found to be 0.66 mM, due to the combination of steepness of the curve, absorbance level and most reproducible results. In these tests, the final concentration of cresol red and DRR was kept constant at 0.05 mM and 12.5 µg/mL, respectively.

*S4. Serial dilution of fluorescein*

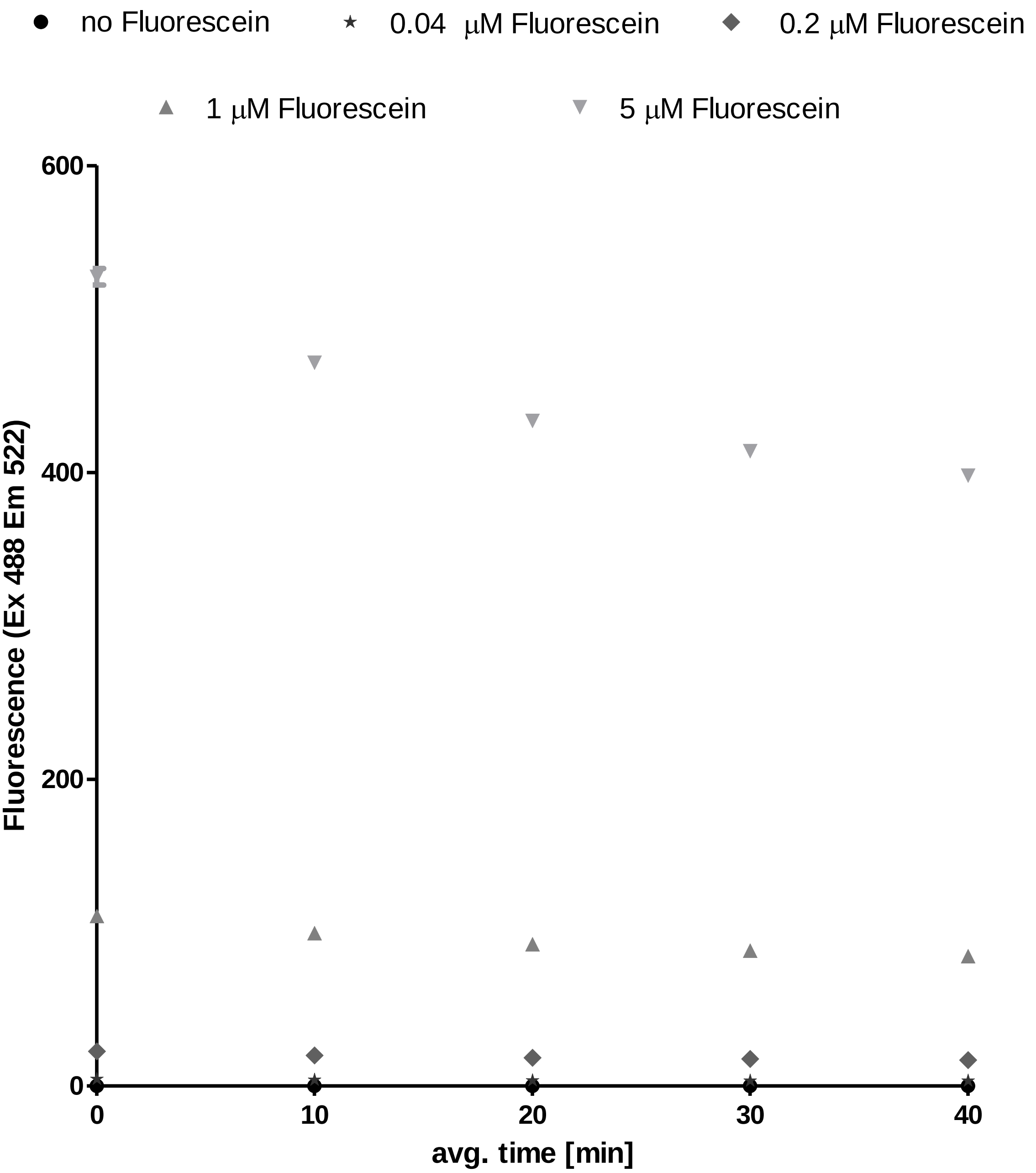


Figure S4. Optimization of the fluorescein concentration in the fluorescein based $PLA_2$ assay. The different assay concentrations tested were 5, 1, 0.2, 0.04, and 0.0 μM fluorescein in presence of 12.5 μg/mL DRR venom. Each measurement point represents the mean of three measurements and the error bars represent SEMs. All concentrations tested gave a similar, concentration-dependent, repeatable and stable fluorescence decrease upon acidification of the assay medium. From these results, a fluorescein assay concentration of 1 μM was chosen. In these tests, the final concentration of phosphatidylcholine and DRR was kept constant at 0.66 mM and 12.5 μg/mL, respectively.
